# Perceptual priors add sensory detail to contextual feedback processing in V1

**DOI:** 10.1101/2023.09.23.559098

**Authors:** Yulia Lazarova, Yingying Huang, Lars Muckli, Lucy S. Petro

## Abstract

How do we develop models of the world? Contextualising ambiguous information with previous experience allows us to form an enriched perception. Contextual information and prior knowledge facilitate perceptual processing, improving our recognition of even distorted or obstructed visual inputs. As a result, neuronal processing elicited by identical sensory inputs varies depending on the context in which we encounter those inputs. This modulation is in line with predictive processing accounts of vision which suggest that the brain uses internal models of the world to predict sensory inputs, with cortical feedback processing in sensory areas encoding beliefs about those inputs. As such, acquiring knowledge should enhance our internal models that we use to resolve sensory ambiguities, and feedback signals should encode more accurate estimates of sensory inputs. We used partially occluded Mooney images, ambiguous two-tone images which are difficult to recognise without prior knowledge of the image content, in behavioural and 3T fMRI experiments to measure if contextual feedback signals in early visual areas are modulated by learning. We show that perceptual priors add sensory detail to contextual feedback processing in early visual areas in response to subsequent presentations of previously ambiguous images.

## Introduction

Context matters. Navigating through a dimly lit room that is familiar is easier than a room you have never been to. Memory and experience provide important context to visual perception. Neuronal responses to sensory signals in V1 depend on the neuron’s response to other, nonsensory, inputs (Gilbert and Li, 2013; Muckli and Petro, 2013). These contextual interactions include cortical feedback inputs from higher visual areas supporting perceptual organisation of scene elements (Roelfsema, 2006), top-down influence from other sensory modalities e.g. auditory (Vetter et al. 2014, 2020), and the encoding of sensory predictions (Alink et al., 2010, Ekman et al., 2017). Therefore, investigating cortical feedback processing is essential for understanding perceptual computations from the perspective of predictive processing frameworks. However, there is much to understand about the neuronal and circuit mechanisms of feedback processing, including how to incorporate mechanisms by which this processing is constrained by our knowledge or updates to perceptual internal models. In turn, investigating how knowledge influences top-down processing is necessary for understanding how cognitive operations depend on the brain’s ability to use predictive internal models to contextualise sensory coding (Larkum, 2013).

Using functional brain imaging (fMRI), we are confronted with a methodological problem. During visual stimulation, contextual feedback signals in V1 are intermixed with sensory signals, so it is impossible to measure these two processing streams in isolation. Previously, we have taken advantage of V1’s retinotopic organisation to isolate non-feedforward signals in the human brain using fMRI. We partially masked a portion of the stimulus and measured brain activity in the region of V1 responding to this occluded region. Brain activity in regions of non-stimulated V1 is related to contextual feedback responses to the surrounding visible scene (Smith and Muckli, 2010, Muckli et al., 2015, Revina et al., 2018, Morgan et al., 2019). To investigate what information contextual feedback signals send to V1 in the case of missing scene information, we tested various models and found that line drawings of missing scene portions created by a group of subjects correlated with brain activity patterns from voxels responding to the same occluded portion of the image (Morgan et al., 2019). The line drawings revealed internal models representing ‘best guesses’ based on world knowledge; they were not created by memory traces as the subjects were never presented with the full images. Further, these line drawings were consistent across subjects. This finding was timely because in spite of established sensory features that drive V1 neurons, we had less understanding of the structure of our brains’ internal models. However, we still do not know how the expectation of scene features in the occluded region depends on our ability to recognise and make sense of the surrounding scene, i.e. on our perceptual knowledge. For example, an object like a partially visible car parked behind a tree can lead to a low-level prediction that is fed back to V1 representing the contours of a likely continuation of lines consistent with Gestalt laws of figures, or representing a more detailed internal model of that particular model of car. We hypothesised that recurrent levels of visual cortex construct internal models of low-level scenes features that are communicated back to V1, that might be independent of recognition. However, given that best estimates of sensory inputs depend on experience, we further hypothesise that perceptual priors add sensory detail to contextual feedback processing.

To investigate the role of knowledge in updating internal models, we chose ambiguous two-tone Mooney images combined with a visual occlusion paradigm. Mooney images are difficult to recognise without prior knowledge (Castelluccia et al., 2017; Imamoglu et al., 2012). We performed a behavioural recognition task to test whether prior knowledge of category improves the recognition of ambiguous images. We then used Mooney images in an occlusion paradigm in a 3T fMRI experiment to test how contextual feedback signals in V1, V2, and V3, are influenced by recognition. Do Mooney images trigger contextual feedback processing based on the local image features (contours of black and white patches) or based on the knowledge that those contours belong to certain objects? We presented the same Mooney images before and after exposing our subjects to the greyscale images from which the Mooney images were created. Does exposure to the greyscale images help our subjects to recognise content during the second presentation of Mooney images? Does knowledge acquired from the greyscale image enrich top-down contextual information? We investigated whether this enrichment of information can be detected in the feedback signal to retinotopic visual areas.

## Methods

### Participants

We recruited participants with normal or corrected-to-normal vision through the Glasgow University School of Psychology and Neuroscience Subject Pool. 20 healthy participants attended the behavioural experiment (7 males; 18-31 years old). We excluded data from one participant due to missing responses on a large number of trials. We recruited an unrelated group of 25 healthy volunteers for the 3T fMRI experiment (10 males; age range: 20 to 37 years, mean age: 25 years). The research was approved by the ethics committee at the University of Glasgow College of Science and Engineering (300180284), and conforms to the World Medical Association Declaration of Helsinki. Volunteers gave written informed consent and received compensation for participation.

### Stimuli

We sourced stimuli from the Flickr (https://www.flickr.com/) and Unsplash (https://unsplash.com/) databases. In the behavioural paradigm, we used images from 4 categories: faces (100 images), landscapes (100 images), animals (100 images) and manmade objects and scenes (50 images). All original images were in greyscale. We used a Python script (Python 2.7) to convert half the images from each of the 3 main categories (faces, animals, and landscapes) and all images from the manmade category to two-tone Mooney images following the methodology described by Imamoglu et al. (2012, Figure 1). We smoothed the greyscale images (sigma levels between 3 and 6, based on responses from 5 volunteers) and thresholded before converting to Mooney images. All images were resized to 500 x 500 pixels. Following the same procedure, we generated six new images for the fMRI paradigm: 2 faces, 2 landscapes and 2 animals. We positioned a grey occluder over the lower right quadrant of each of the 6 Mooney images. The resulting occluded two-tone images and their corresponding non-occluded greyscale versions formed the final 12 stimuli used in the fMRI paradigm. We programmed the stimulation code using Presentation® software (Version 18.0, Neurobehavioral Systems, Inc., Berkeley, CA, www.neurobs.com).

**Figure 1.**
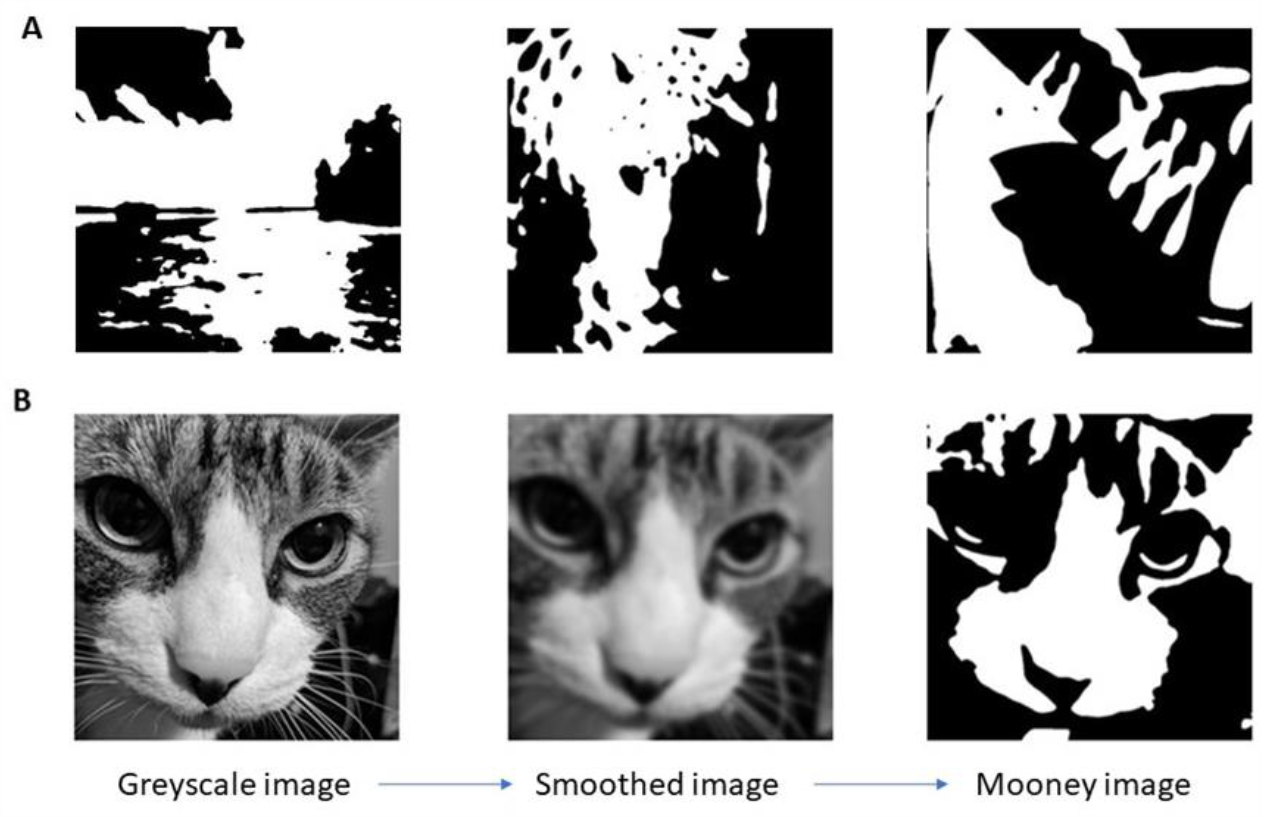
Experimental stimuli. (A) Example two-tone images from different categories: landscapes, animals, and manmade objects. From left to right: beach landscape, leopard, shoe. (B) Mooney images generation procedure: greyscale images were downloaded from the Flickr and Unsplash databases. To create the Mooney stimuli, we smoothed and thresholded each image following the procedure described by Imamoglu et al. (2012).

### Behavioural study design

In the behavioural experiment, we wanted to test whether prior knowledge of category improves the recognition of ambiguous images. The experiment consisted of 4 runs (Figure 2). In run 1, participants saw 150 Mooney images presented in a structured category order: face, landscape, animal. In run 2, they saw 150 novel greyscale images presented in the same category order. In run 3, we presented 150 Mooney images in the same category order as runs 1 and 2. To control for the effect of repeated exposure to the same stimuli, 50% of the images presented in run 3 were already seen in run 1 and the remaining 50% were novel images. This percentage was balanced across categories and the novel and previously seen images were presented in a semi-random order, keeping the category order consistent in runs 2 and 3. In run 4, participants saw 200 two-tone images, presented in a random order. 150 of the images used in run 4 were from the three known categories, 50% of which were already seen once before and the remaining 50% were novel two-tone images. Run 4 included an additional category of 50 images depicting man-made objects which were shuffled amongst the other three categories. Runs 1, 2 and 3 were pseudo-randomised within blocks, to ensure category order was maintained and images within blocks were randomised across subjects, so each subject saw a different combination of images.

**Figure 2.**
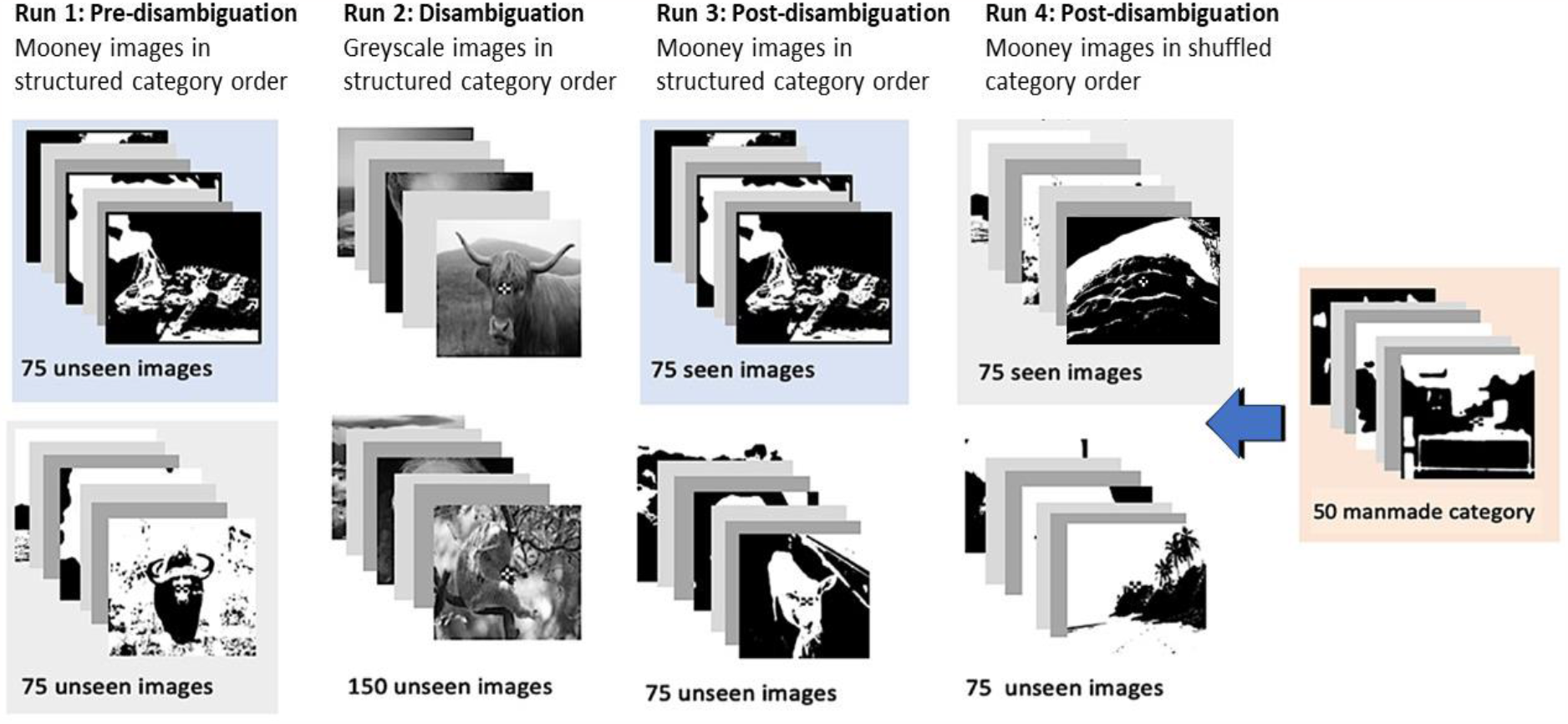
Experimental paradigm used in the behavioural study. There were 4 experimental runs: run 1 consisted of 150 two-tone images; run 2 consisted of 150 non-identical greyscales from the same categories as the images in run 1, and presented in the same order; run 3 presented 75 Mooney images seen in run 1 and 75 new Mooney images ; run 4 presented remaining 75 Mooney images from run 1 and 75 unseen before Mooney images shuffled together with the 50 images of the manmade category and presented in a random order.

### Behavioural study procedure

Participants were seated in front of a 23-inch Dell OptiPlex 9030 Monitor (1920 × 1080; 59 Hz) with their head resting on a chinrest positioned 60cm from the screen and set to a 32cm height from the desk. Participants were given no information about the categories of the images or the presentation order. Images were presented on the screen for 1 second. Each two-tone image was followed by 3 questions (Figure 3): whether they could recognise what was depicted in the image (keyboard button y = yes and n = no); if they gave a positive response, they were asked to type in 1 to 3 words what they saw in the image, alternatively they were asked to guess what was in the image; finally they indicated their confidence in the answer on a continuous scale from 1 (I am not very confident at all) to 4 (I am very confident). At the start of run 2, a message on the screen instructed participants that they were about to see the greyscale images and asked them to press 1 when they saw a face, 2 for an animal and 3 for a landscape. They responded when prompted with a fixation cross colour change ensuring they attended to the stimuli rather than rely on temporal regularity. During this disambiguation phase, they gained knowledge of the regularity of category order but not of the exact greyscale versions of the two-tone images seen before. Participants were given breaks after each run. The experiment lasted approximately 90 minutes.

**Figure 3.**
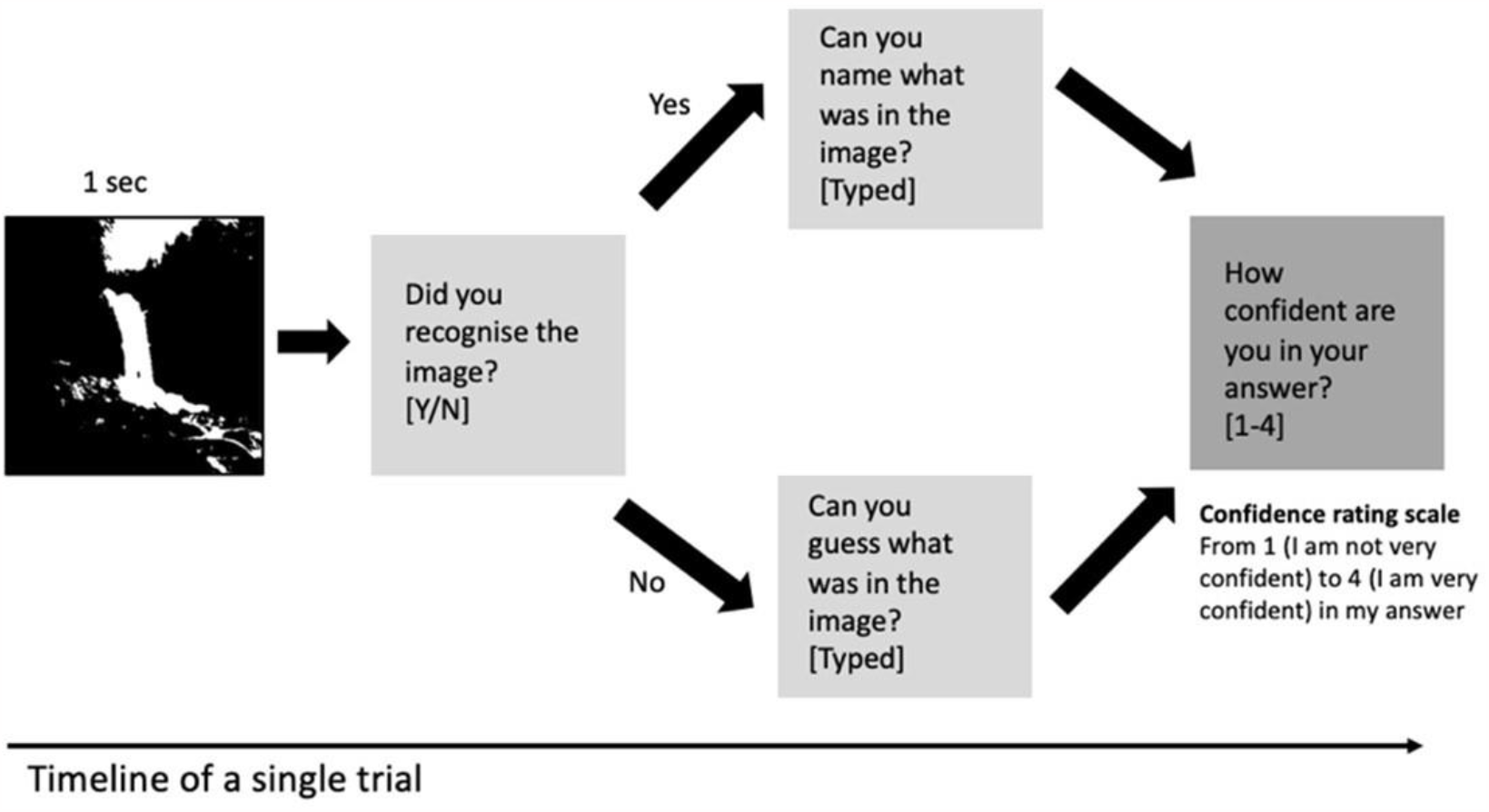
The timeline of a Mooney image trial in the behavioural study. The two-tone image was presented for 1 sec. Then the participants were asked to confirm whether they recognised what is depicted in the image (yes/no). Upon a positive response they were asked to type what they saw in the image with 1-3 words, upon a negative response they were asked to guess what the image depicted. Finally, they rated the confidence in their response on the scale from 1 (low confidence) to 4 (high confidence).

### fMRI study experimental design

We followed the canonical Mooney image paradigm of 3 phases: pre-disambiguation, when novel occluded two-tone images are presented; disambiguation, presenting the full greyscale versions of the images seen before; post-disambiguation, a repetition of the same Mooney images from phase 1 (Figure 4). We used a block design where each image was presented for 6 seconds followed by a 6-second intertrial interval. During the 6-second on interval the image was flashed on the screen at a 5Hz, to give an optimal signal-to-noise ratio of the BOLD response to complex visual stimuli (Kay et al., 2008). The images appeared in semi-randomised order, they were shuffled within categories but presented in a fixed category order: landscape, face, animal. Each individual image was repeated 8 times in a run. We recorded 2 runs of functional data pre-disambiguation, and 2 runs post disambiguation. To be able to functionally locate the non-stimulated patch of visual cortex corresponding to the occluded region, we included two types of mapping trials spanning around the border of the occluded region and covering the rest of the occluded region (Figure 4B). For the stimulation during the retinotopic mapping, we combined a polar angle wedge and an eccentricity ring (Figure 4C).

**Figure 4.**
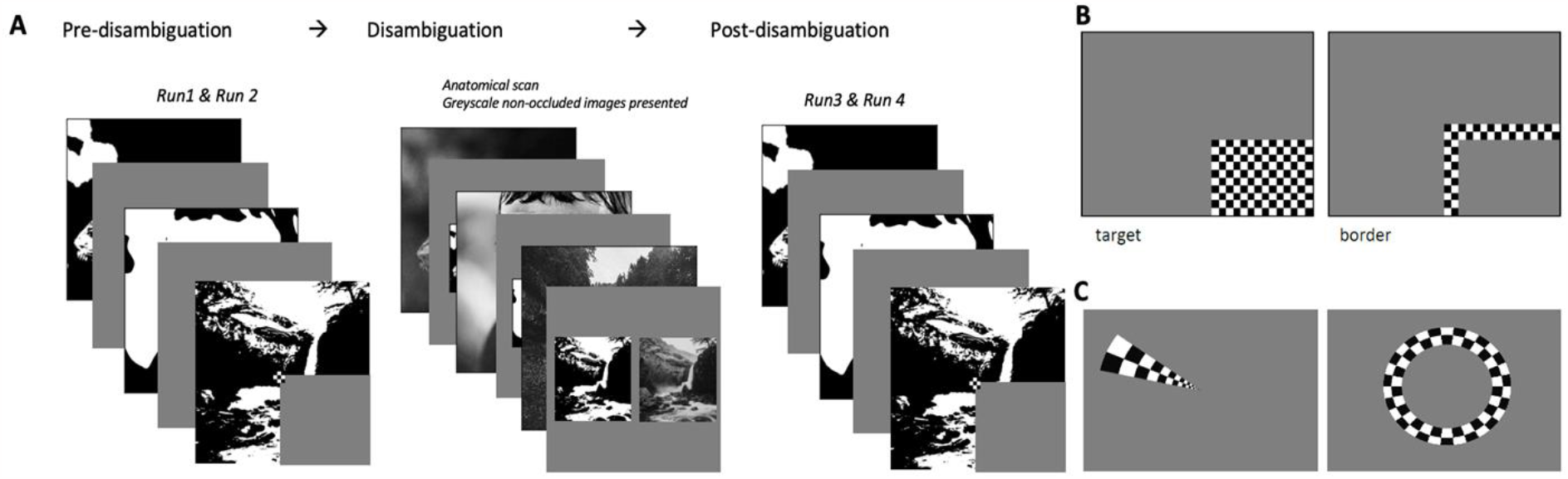
Experimental paradigm used in the fMRI experiment. A) Study paradigm and example stimuli. In runs 1 and 2 (pre-disambiguation phase) participants saw the occluded Mooney images. While the anatomical scan was recorded we performed the disambiguation with the greyscale versions of the same images. In runs 3 and 4 (post-disambiguation phase) we repeated the Mooney images from run 1. B) Mapping trials used to locate the voxels responding to the occluded region of the visual field. C) Stimulation used for retinotopic mapping, comprising polar angle and eccentricity mapping.

### fMRI procedure

The stimuli were displayed on a rear-projection screen using a projector system (1024 x 768 resolution, 60 Hz refresh rate). A centralised fixation checkerboard (9 x 9 pixels) marked the centre of the scene images. Participants were not given category information about the images prior to the experiment. Participants were instructed to keep their eyes fixated on the central fixation cross. For each Mooney image presented on the screen they were asked to respond by pressing one of two buttons (yes and no) to indicate if they recognised what was in the currently presented image. They were instructed to give a yes response only if they could tell what the image depicted and not to indicate that they recognise the image from having seen it in a previous trial. They were prompted for a response by a colour flicker on the fixation cross. We recorded 2 functional runs of occluded Mooney images pre-disambiguation. Following the two runs, we recorded a structural image of the brain while presenting the participants with the disambiguation images. The full greyscale equivalents of the Mooney images seen previously were presented consecutively on full screen then side by side with the corresponding non-occluded Mooney version of the image. During this phase, participants were allowed to freely explore the images without the need to fixate their gaze and to remember each image in detail. Post-disambiguation, we recorded two more functional runs repeating the main Mooney image recognition task. We concluded the scanning session by recording a combined polar and eccentricity retinotopic map (Figure 4C), which we later used to functionally identify our regions of interest.

### fMRI acquisition

We recorded MRI data at the Centre of Cognitive Neuroimaging in the University of Glasgow. We used a 3T Tim Trio MRI system (Siemens, Erlangen, Germany) equipped with a 32-channel head coil to record the T1-weighted anatomical and echo-planar (EPI) images. For the four experimental runs we used an EPI sequence to acquire partial brain volumes aligned to maximise full coverage of the early visual cortex (18 slices; voxel size: 3 mm, isotropic; 0.3 mm inter-slice gap; TR = 1000 ms; TE = 30 ms; matrix size = 70×70; FOV = 210 mm, 817 vols, flip angle of 62 deg, PAT mode GRAPPA (accel factor of 2), echo spacing of 0.49 ms, online motion correction and a bandwidth of 2464 Hz/Px). We acquired a high-resolution anatomical scan (3D MPRAGE, voxel size: 1 mm, isotropic, 192 vols). The acquisition parameters of the retinotopic functional run were as follows: 38 slices, voxel size: 3 mm, isotropic; 0.3 mm interslice gap; TR = 1000 ms; TE = 26 ms; matrix size = 74×74; FOV = 222mm, 796 volumes, Multiband acceleration factor of 2, flip angle of 60 deg, PAT mode GRAPPA (accel factor of 2), echo spacing 0.48 ms, online motion correction and a bandwidth of 2702 Hz/Px, EPI factor 74.

### fMRI data pre-processing

We used BrainVoyager 22.0 (Brain Innovation, Maastricht, The Netherlands) (Goebel et. al., 2006) for preprocessing the fMRI data. On each functional run we applied slice scan time correction, using cubic spline interpolation, and 3D motion correction with Trilinear/Sinc interpolation. To account for motion within as well as between runs we aligned all functional runs to the first volume of run 3 which was closest in time to the anatomical scan. Additionally, we applied a temporal high-pass filter with GLM-Fourier basis set at 7 cycles, and linear detrending. We co-registered this data with the anatomical data and brought it into Talairach space.

### ROI definition

We applied phase encoded retinotopic mapping using cross-correlation analysis on the retinotopic mapping data to functionally define early visual areas V1, V2, and V3. Additionally, we performed a general linear model (GLM) on the mapping trials from the main functional runs, contrasting the target and the border mapping trials. To eliminate as much as possible the contribution from lateral interaction effects, we selected only voxels which responded to the target but not the border region. The identified patches of voxels within V1, V2, and V3, corresponding to the occluded section of the visual field. This resulted in 3 non-stimulated regions (V1, V2, and V3, see examples in Figure 5) and three stimulated regions (within V1, V2, and V3) which were our main regions of interest for subsequent analyses. Non-stimulated regions refer to regions receiving uninformative feedforward information, responding only to the grey screen. Therefore stimulus-specific brain activity in this region is related mainly to feedback processing, potentially with some contribution from lateral inputs though we minimise this with our ‘surround’ mapping checkerboards. Stimulated regions received sensory stimulation from Mooney images, and also contextual feedback signals from higher areas, and lateral inputs. For simplicity, we refer to feedforward and feedback processing conditions, in stimulated and non-stimulated regions respectively.

**Figure 5.**
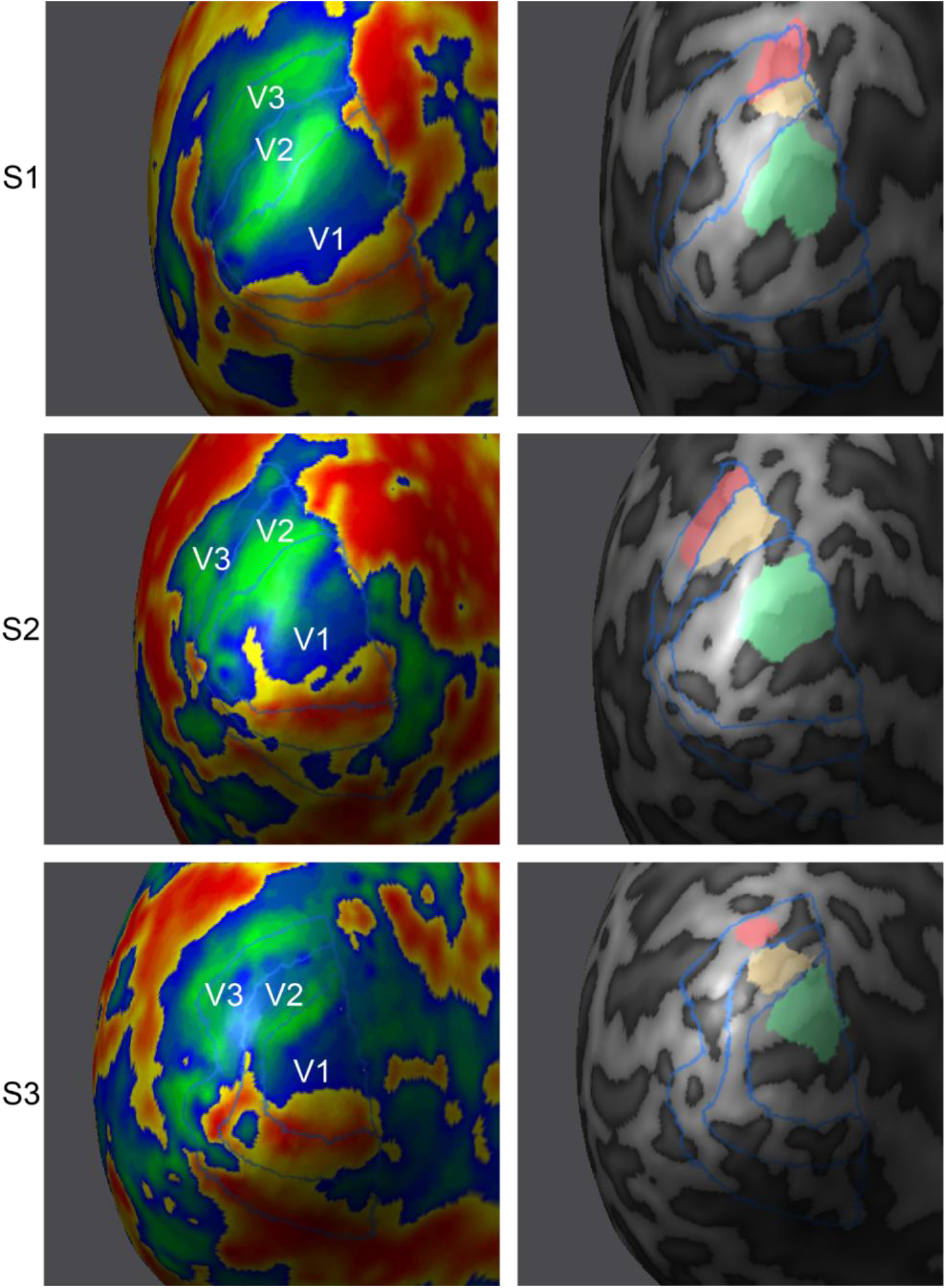
Non-stimulated regions localised in the early visual cortex of the left hemisphere. Here, we present three subjects’ left hemisphere inflated meshes. The left column is the result of retinotopic mapping and shows borders between early visual areas. The right column shows the retinotopic location of the white occluder in the stimulus image, for V1 (green), V2 (yellow), V3 (pink). These ‘non-stimulated’ regions receive feedback (and lateral) processing, and uninformative feedforward information.

### Behavioural data analyses

We analysed the behavioural data (N=19) using R (version 3.6.0, using package lme4 by Bates et al., 2015). To investigate the effect of priming on interpreting ambiguous visual information, we compared image recognition rates prior to disambiguation (run 1) to those post-disambiguation: in a predictable image order (run 3) and unpredictable shuffled order (run 4). To control for confounding effects of repeated exposure to the same Mooney image, we compared the responses given to seen images as opposed to novel ones in runs 3 and 4. To control for the effect of learning we compared responses between the three categories experienced previously to the novel category of manmade objects in run 4. We used the ‘yes’ and ‘no’ response count as a measure of recognition rate. We applied a generalised linear mixed model (GLMM) test, which accounted for repeated-measures dependencies in a within-subjects and within-items design in all cases, with full random effects structure.

### fMRI analysis - SVM classification

We performed multivoxel pattern analysis (MVPA) training a linear Support Vector Machine (SVM) algorithm to decode the differences in cortical feedback activity in early visual areas in response to Mooney images. First, we extracted voxel activation patterns of each trial extracted from each ROI and then z-scored the parameter estimates (β values). We then split the data from each phase: pre- and post-disambiguation into a training set (50% of the data) and a testing set (the remaining 50%). We employed this cross-validation approach of training and testing the SVM classifier on separate data sets to avoid overfitting of the model. For each of the previously defined feedforward processing (stimulated) and feedback processing (non-stimulated) ROIs, we obtained a pre- and post-disambiguation classification accuracy for each of the six images. We applied permutation testing of the SVM classification to determine the significance of individual participants. The permutation was performed by shuffling the data labels in the training set and leaving the labels of the testing set intact. Performing 1000 repetitions of this procedure yielded a distribution around chance-level (decoding at 50% accuracy). We compared the observed classification value to this distribution to determine the significance of the classifier’s ability to successfully decode the individual images.

### fMRI Classification analysis - RSA

To compare the neuronal representations across the different phases of recognition, we compared the similarities of the neural responses to individual images in pre- and post-disambiguation phases. We applied a representational similarity analysis (RSA), aiming to increase the sensitivity to the difference in patterns of responses by exploiting the continuous nature of the RSA score (Kriegeskorte et al., 2008). We were interested in changes in the image representations introduced by the disambiguation phase. We computed dissimilarity matrices for each ROI comparing images within as well as across categories for each of the redefined ROIs. In this process responses to every single image were compared to every other image in the set of stimuli. Representational dissimilarity matrices (RDMs) were calculated by 1 minus linear Pearson correlation coefficient (1-r). Further, we also conducted statistical analysis (t-test) for group mean dissimilarity between pre- and post-disambiguation phases in each ROI.

## Results

### Behavioural experiment

We compared recognition rates pre- and post-disambiguation to understand whether category information was enough to improve recognition rates even in the absence of image-specific information. To measure this priming effect, we used a GLMM analysis. We compared responses to the three primed categories (animals, faces, and landscapes) pre-disambiguation (run 1) to post-disambiguation (runs 3 and 4, excluding the manmade image category). We normalised scores for runs 3 and 4 to control for the difference in the number of image presentations between pre- (run 1) and post-disambiguation phase (runs 3 and 4).

We found increased recognition scores following the disambiguation phase (Figure 6A). The GLMM analysis revealed a main effect of priming: LRχ2 = 20.16, p = 7.11e-06. To test for the effect of repeated exposure to the same Mooney image, we compared responses to seen and unseen images that were presented post disambiguation (Figure 6B). The model containing the repetition effect did not explain the observed data better than the model that does not contain the repetition effect (main effect of repetition: LRχ2 = 0.04e-01, p = 0.95), therefore the repetition effect did not contribute significantly to the increase in recognition rates we observed post disambiguation. To further ensure that the increase in recognition rates is due to the predictable category order we added to our model a term for the image presentation order, which further improved our model’s fit to the data (main effect of order: LRχ2 = 7.59, p = 0.01; see Figure 6C). Finally, we were interested to see if the general expectation of one of the three familiar categories was enough to increase recognition even in absence of structured presentation order. We compared responses to images from the primed categories (landscape, face, animal) and images to the unprimed category (manmade) in run 4 (see Figure 6D). We found higher recognition performance (main effect of primed category: LRχ2 = 60.31, p = 8.10^e-15^) for the primed categories compared to the newly introduced one.

**Figure 6.**
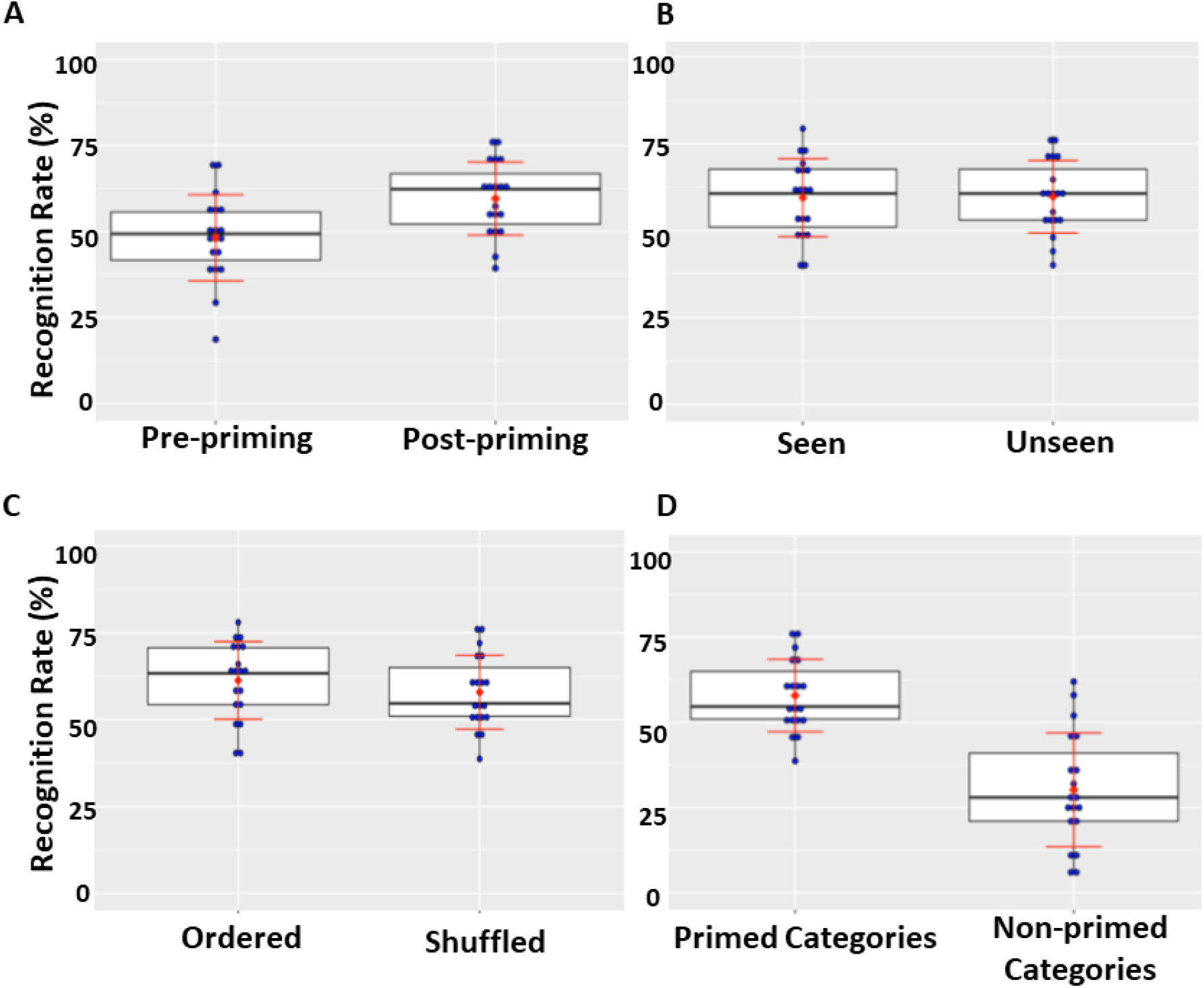
Percentage of Mooney images recognised. The red dot in each plot marks the mean percentage of recognised images across individuals. Individual subjects’ scores are marked in blue. The error bars indicate the standard deviation. (A) Comparing recognition rates between pre- (run 1) and post- (run 3 and run 4) disambiguation. (B) All data shown here is from the post-disambiguation phase (run 3 and 4) - percentage of seen (repeated from run 1) and unseen images recognised post-priming runs 3 and 4. (C) Percentage of Ordered (run 3) and shuffled (run 4) images recognised post-priming. (D) Recognition rates between primed categories (face, animal, and landscape) and un-primed category (manmade) in run 4.

### fMRI - MVPA Classification

We applied SVM classification on data from each phase: before and after presenting participants with the greyscale versions of the images. The classifier could successfully dissociate between the images above chance level across all ROIs. This was the case when training and testing the classifier on data from feedforward activations as well as feedback activations (Table 1, Figure 7). A GLMM analysis revealed a main effect of the phase in nonoccluded regions of V2 and V3 but not in V1 (V1: t=-1.47, df=748, p=0.141; V2: t=-2.36, df=748, p=0.018; V3: t=-2.00, df=748, p=0.046). For the occluded regions we saw a significant main effect of the phase in V1 and V2 (V1: t=-6.08, df=748, p=1.946^e-09^; V2: t=-4.58, df=748, p=5.583^e-06^; V3: t=-2.13, df=748, p=0.034).

**Table 1.** Summary statistics and significance results for the classifier decoding accuracy. The classifier could successfully decode the images from each other above chance level (50%) based on activations from both feedforward (FF) and feedback (FB) activations from voxels in each roi: V1, V2 and V3. This was true for data recorded before (PRE) and after (POST) disambiguation. The mean values and the 95% confidence intervals reported are in %correct classification.

| ROI | FF/FB | Phase | t | df | mean | CI (95%) | p-value |
| --- | --- | --- | --- | --- | --- | --- | --- |
| V1 | FF | PRE | 63.278 | 374 | 0.89 | 0.878 - 0.903 | <0.001 |
| V1 | FF | POST | 54.246 | 374 | 0.88 | 0.866 - 0.894 | <0.001 |
| V1 | FB | PRE | 37.339 | 374 | 0.78 | 0.767 - 0.796 | <0.001 |
| V1 | FB | POST | 26.517 | 374 | 0.73 | 0.710 - 0.743 | <0.001 |
| V2 | FF | PRE | 54.167 | 374 | 0.87 | 0.857 - 0.884 | <0.001 |
| V2 | FF | POST | 47.368 | 374 | 0.85 | 0.836 - 0.865 | <0.001 |
| V2 | FB | PRE | 21.86 | 374 | 0.65 | 0.639 - 0.667 | <0.001 |
| V2 | FB | POST | 14.764 | 374 | 0.61 | 0.598 - 0.628 | <0.001 |
| V3 | FF | PRE | 47.844 | 374 | 0.84 | 0.830 - 0.858 | <0.001 |
| V3 | FF | POST | 40.835 | 374 | 0.83 | 0.810 - 0.842 | <0.001 |
| V3 | FB | PRE | 14.896 | 374 | 0.60 | 0.589 - 0.617 | <0.001 |
| V3 | FB | POST | 12.358 | 374 | 0.58 | 0.57 - 0.598 | <0.001 |

**Figure 7.**
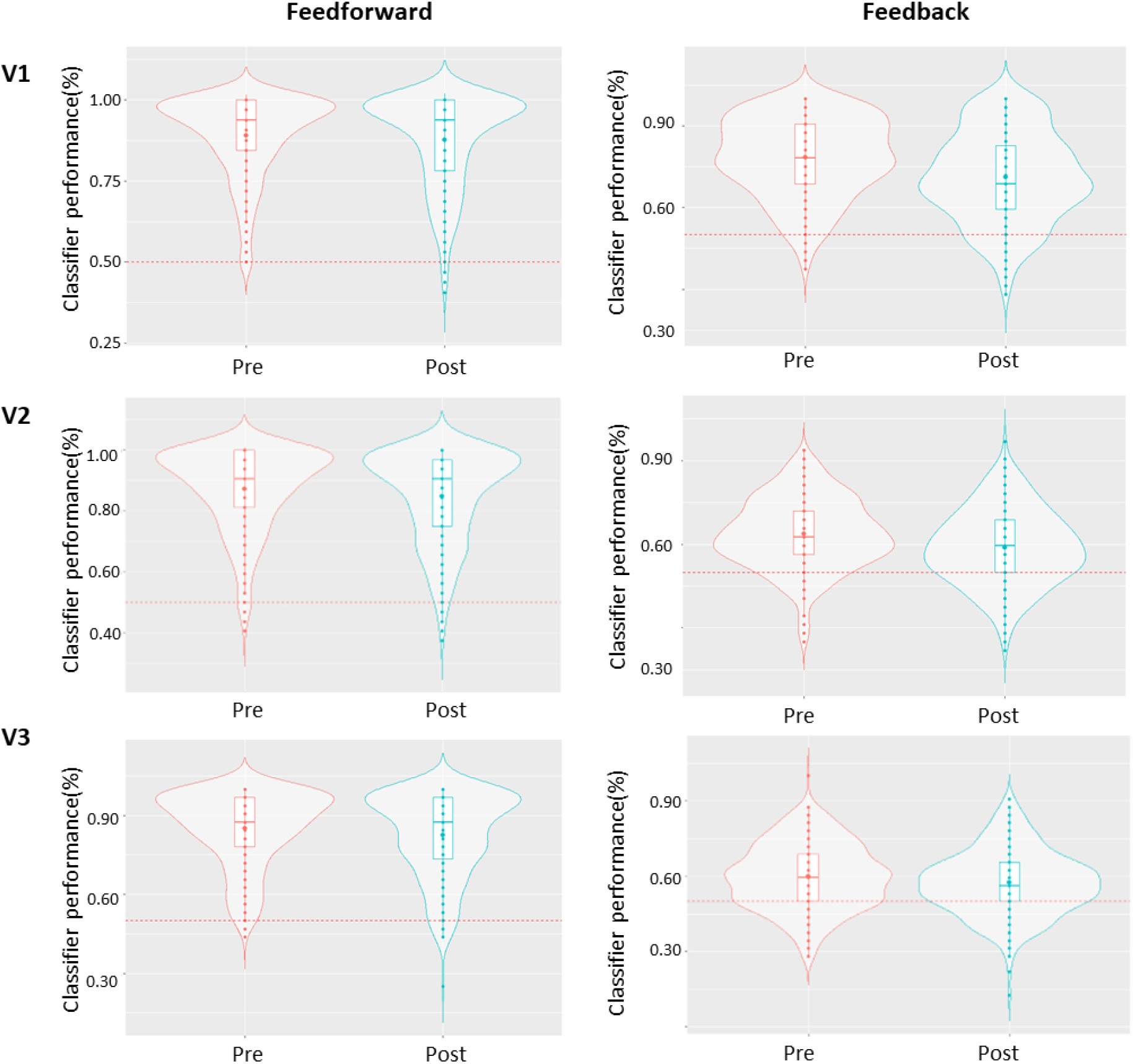
Classification accuracy pre- and post-disambiguation. Classification accuracy before (red) and after (green) presenting the greyscale versions of the images. The left column of plots shows results for non-occluded (feedforward) sections of V1, V2, and V3 (top to bottom), the right column of plots represents results for occluded (feedback) regions within the respective ROI. Each boxplot represents the median and the interquartile range of the data and the violin plots represent the data distribution. The red dashed line indicates the chance level (50%) of decoding accuracy. The classifier performed above chance level in all conditions and for each ROI (for confidence intervals and significance please refer to table 1).

### RSA

To further investigate the effect of priming on forming internal representations of the ambiguous images we performed representational dissimilarity analysis. Individual comparisons across all images are presented in the RDMs in Figure 8 and Figure 9 along with the overall increase in dissimilarity following the disambiguation phase. We saw a significant increase in dissimilarity between individual images in all ROIs for both feedforward activations, between pre and post disambiguation phases (Figure 8) (V1: t= -11.05, df=14, p<0.001; V2: t= -12.87, df=14, p<0.001; V3: t= -13.82, df=14, p<0.001) as well as feedback activations (Figure 9) (V1: t= -9.67, df=14, p<0.001; V2: t= -12.38, df=14, p<0.001; V3: t= - 10.07, df=14, p<0.001).

**Figure 8.**
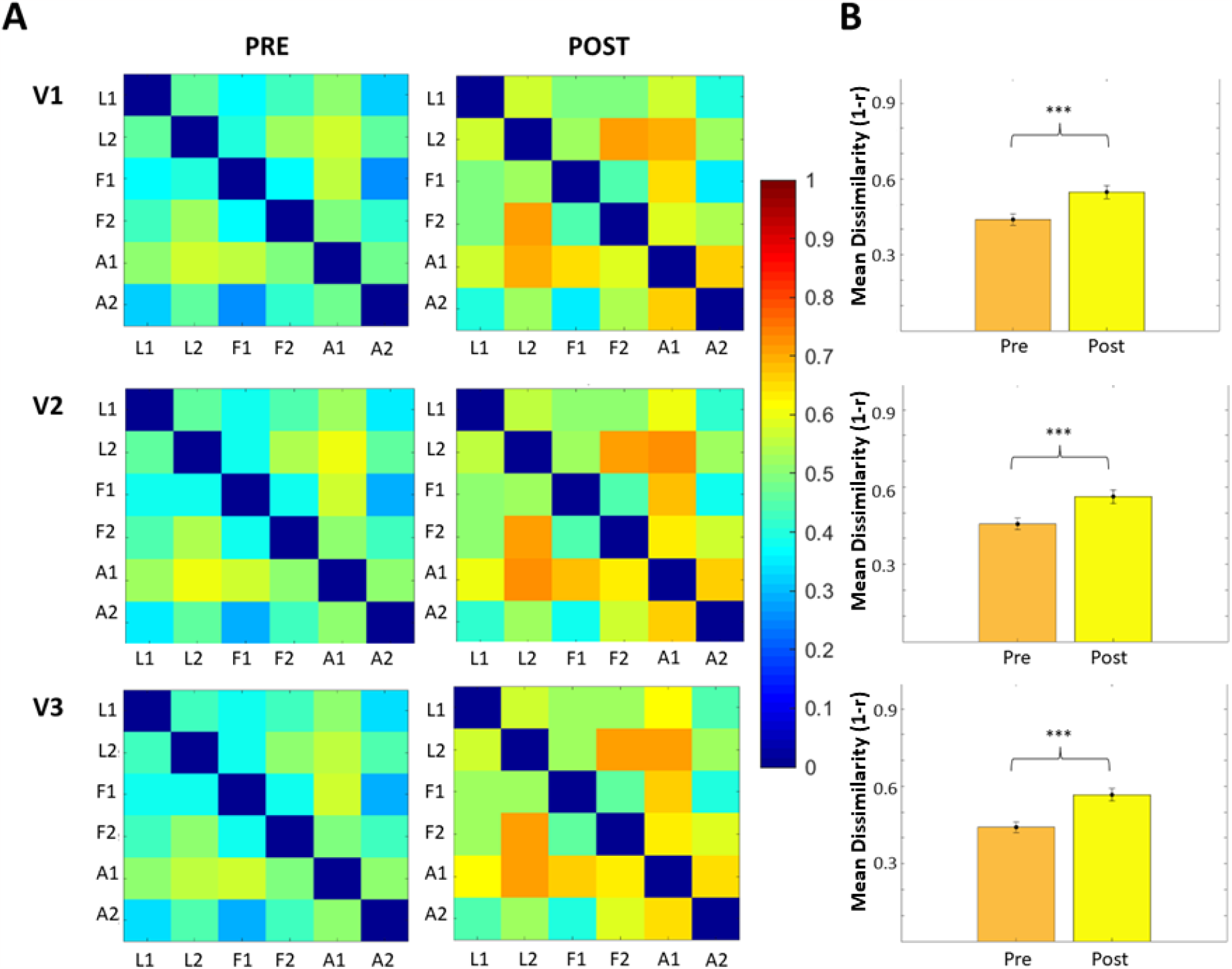
Dissimilarity matrices and mean dissimilarity for feedforward activations in each of the three ROIs. (A) The two columns of RDMs show mean dissimilarity between individual images (Landscape image 1 = L1, Landscape image 2 = L2, Face image 1 = F1, Face image 2 = F2, Animal image 1=A1, Animal image 2= A2). The left and the column of RDMs show results before disambiguation (PRE) and after disambiguation respectively (POST). Warmer colours on the scale indicate higher dissimilarity between activations elicited by the images, cooler tones indicate lower dissimilarity. (B) Bar graphs representing the mean dissimilarity between images before (orange) and after (yellow) presenting the greyscale images. The dissimilarity is calculated by the 1 – r (r is the Pearson linear correlation coefficient). The error bars indicate the 95% confidence interval around the mean. The *** denote the p<0.001 significance of the difference between the two conditions. The pre- and post- RDMs along with the bar graph with mean values are presented for each one of the ROIs V1 on the first row, V2 on the second row and V3 at the bottom of the figure.

**Figure 9.**
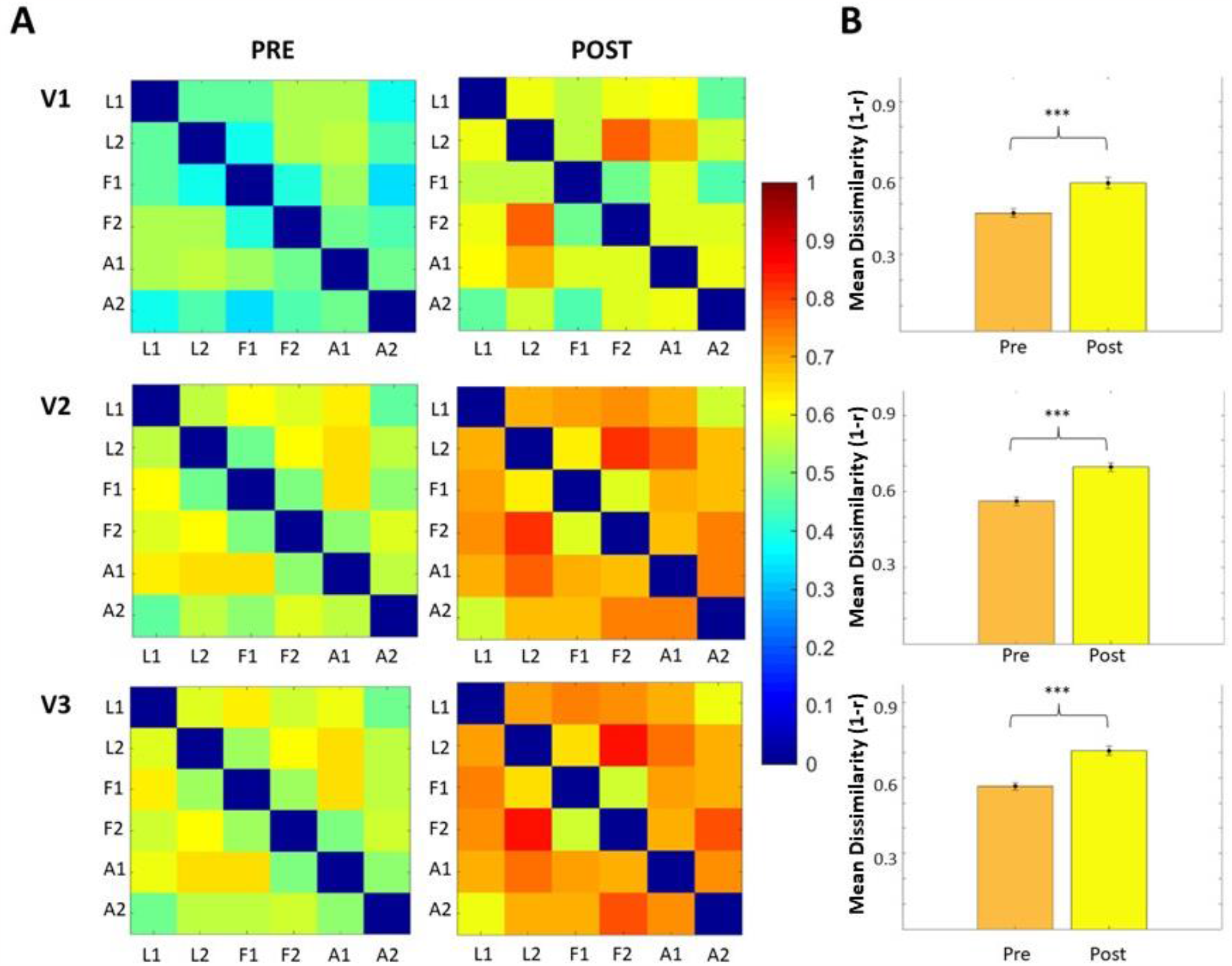
Dissimilarity matrices and mean dissimilarity for feedback activations in each of the three ROIs. (A) The two columns of RDMs show mean dissimilarity between individual images (Landscape image 1 = L1, Landscape image 2 = L2, Face image 1 = F1, Face image 2 = F2, Animal image 1=A1, Animal image 2= A2). The left and the column of RDMs show results before disambiguation (PRE) and after disambiguation respectively (POST). Warmer colours on the scale indicate higher dissimilarity between activations elicited by the images, cooler tones indicate lower dissimilarity. (B) Bar graphs representing the mean dissimilarity between images before (orange) and after (yellow) presenting the greyscale images. The dissimilarity is calculated by the 1 – r (r is the Pearson linear correlation coefficient). The error bars indicate the 95% confidence interval around the mean. The *** denote the p<0.001 significance of the difference between the two conditions. The pre- and post- RDMs along with the bar graph with mean values are presented for each one of the ROIs V1 on the first row, V2 on the second row and V3 at the bottom of the figure.

## Discussion

As the saying goes, ‘you never step in the same river twice’. We set out to test how the cortex contextualises new information with previously acquired knowledge, and if this information is available even at the level of primary visual cortex. Neuronal responses in the primary visual cortex are influenced by contextual feedback processing. We investigated how contextual feedback processing is altered as a function of prior knowledge. We combined ambiguous two-tone Mooney images with a partial visual occlusion paradigm and 3T fMRI to measure how contextual feedback signals in response to Mooney images are influenced by prior knowledge of the full greyscale versions. The dissimilarity between contextual feedback representations of ambiguous images was greater after the visual system had been presented with the full contrast greyscale versions of the same images. However, we also observed contextual feedback responses in the pre-disambiguation phase related to the two-tone black and white images, prior to exposure to the greyscale source. We discuss these findings in relation to the updating of perceptual internal models.

An increase in knowledge about an ambiguous visual stimulus translates to a more detailed neuronal representation of that stimulus (Gorlin et al., 2012; van Loon et al., 2016), around 500 ms post-stimulus (Flounders et al., 2019). Van Loon et al., administered ketamine, which blocks NMDA receptors, and showed that Mooney images are processed more similarly in early visual areas, suggesting that recognition is impaired by interrupting feedback to early vision. Here, we build on these findings by explicitly separating feedforward sensory processing from feedback processing. Previously, we have shown contextual feedback responses in V1 to natural scene images where the visual context influencing V1 contained higher level visual features (Smith and Muckli, 2010, Muckli et al., 2015, Revina et al., 2018, Morgan et al., 2019). In the current study, we were able to decode Mooney images prior to exposure to the greyscale source image, showing that a more rudimentary visual context modulates V1, potentially coding for contours or low frequency representations of object outlines. This feedback processing might relate to recurrent levels of the visual cortex constructing internal models of lower level scenes features that are sent back to V1 regardless of the image being interpretable or not, and independent from higher level top-down factors.

However, we also found that during this pre-disambiguation phase, prior to greyscale image exposure, contextual feedback neuronal representations were more similar to each other compared to after seeing the full versions of the images. That is, the dissimilarity between contextual feedback representations of ambiguous images was greater after the visual system had been presented with the full contrast greyscale versions of the same images. This could relate to a feedback information code that in the later phase now includes more accurate topdown estimates of the degraded images, based on our knowledge of the features underlying the degraded versions. It is possible that updated internal perceptual models encoded more detailed expected sensory inputs, even when presented with ambiguous images. We speculate that more detailed perceptual priors could be acquired via a one-shot learning type of mechanism in which subjects rapidly learned to associate Mooney images in the post-disambiguation phase to their greyscale counterparts, reducing uncertainty (Lee et al., 2015). Though, we do not intend to distinguish between knowledge and expectation, and it remains to be understood how the knowledge gleaned from seeing the full image interacts with predictive neuronal processing shaping subjects’ expectations of the second presentation of the Mooney image during the post-disambiguation phase, or which cortical stages are involved. The successful decoding of individual images prior to disambiguation indicates that simple low-level properties are driving contextual feedback. Our previous research provided evidence that feedback-activity in occluded visual scenes (we presented 24 partially occluded images of 6 different categories) can be best modelled as mental line drawings of the missing information (Morgan et al. 2019). Here we show that mental line drawings that create decodable activity are driven by the light dark boundaries in the surrounding image without necessarily containing an internal model of the meaning of the surrounding visual scene, and without recognisable objects and other context.

Our findings also suggest that when faced with ambiguous input, the brain automatically associates the current experience with a previously acquired memory to form a perception. This is also evident even from the high proportion of reported subjective image recognition rates prior to disambiguation (Figure 10). Upon debriefing the participants, we learned that they often formed an interpretation of the ambiguous image even prior to seeing its corresponding greyscale version. This initial perception was different from the actual content of the ambiguous image. The majority of participants reported that this initial image was erased from their minds once they saw the original greyscale images. However, for some participants, it persisted even after the disambiguation phase. As a result, the subsequent representation of the ambiguous image might have elicited a competing perception which could account for the decrease in classification performance. In line with this observation, van Dam et al. (2010) reported that prior exposure to ambiguous stimuli can negatively impact the effect of training on the perception of bistable stimuli.

**Figure 10.**
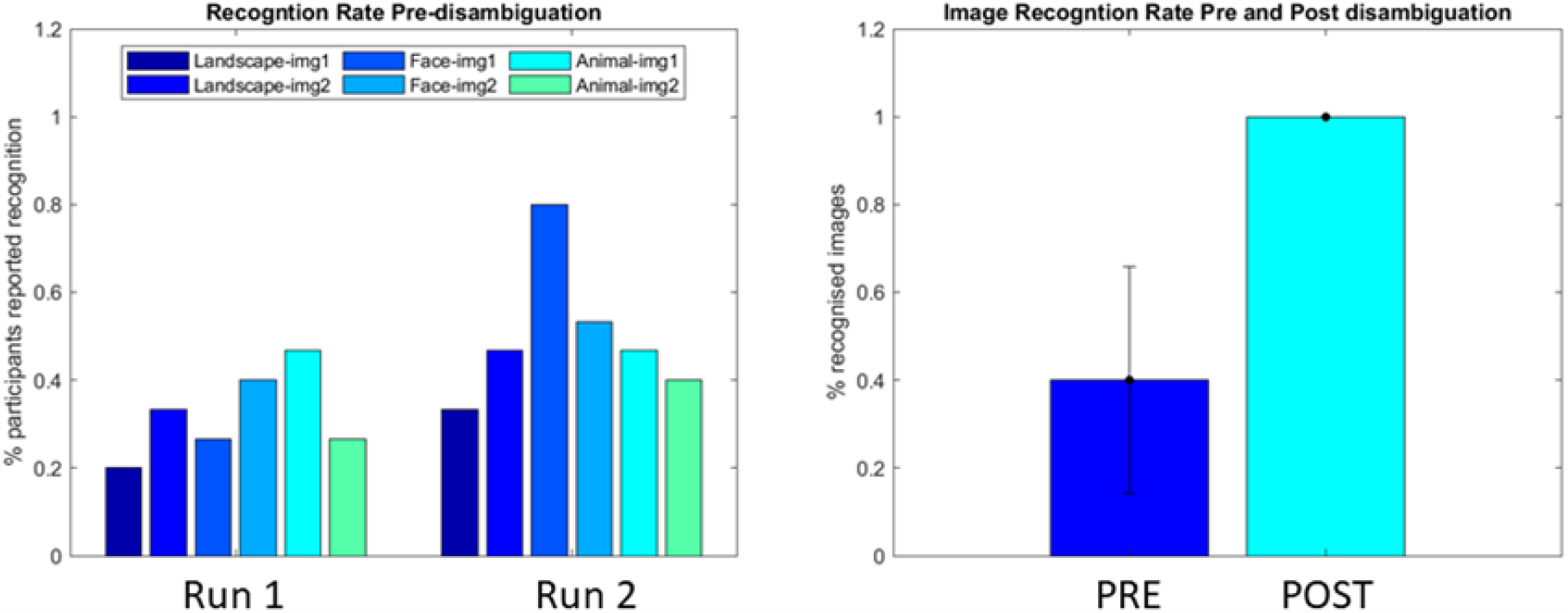
Subjective recognition rate - behavioural data recorded during fMRI scanning. The left panel illustrates the percentage of people who reported they can recognise what is in the image across the two runs prior to disambiguation. The response rates are presented for each individual image. The right panel compares the recognition rate before and after disambiguation averaged across all images.

Our study focuses on the flexible adaptation to context in the adult brain. Here, exposure to a grey-scale image provides a rapid learned context for subsequent exposure to previously ambiguous Mooney images, building the perceptual repertoire over time. Children are typically poor at recognising Mooney images, even when viewing the greyscale counterpart images (Yoon et al., 2007). On a functional level, this finding might relate to not yet having formed rich internal models used to optimise predictions of ambiguous sensory inputs. Animal models of contextual feedback processing in early visual cortex show that feedback information is relevant for recognising objects on backgrounds (Kirchberger et al., 2023). It is possible that in children, contextual feedback processing in response to Mooney images is less informative about the image content than would be so in a more developed or experienced brain, and that enriching contextual feedback is a general hallmark of visual cognitive development. In line with this, Berkes et al., (2011) revealed that primary visual cortical circuits refine internal models to the statistical structure of natural scenes over developmental phases in ferrets.

Lastly, the results from our behavioural study showed that priming with non-identical greyscale training improves the recognition of Mooney images. This priming effect was specific to category-order of images, not repetition of images, and improves recognition even without sequence information. Compared to other informative priming clues such as semantic priming (Samaha et al., 2018), our behavioural experiment demonstrated that even if the available information is limited (e.g., white and black ambiguous category information), participants can still get a priming effect on recognition based on the category-order knowledge. This might indicate the brain extracts information even from impoverished sensory input, to generate predictive templates that generalise across categories of sensory inputs.

Prior experiences and acquisition of knowledge profusely influence our perception. The role of top-down processing carried by cortical feedback signals is widely accepted but we were lacking human brain imaging evidence that feedback signals are modulated by the updating of internal models. Here we show that perceptual priors add sensory detail to contextual feedback processing in V1 to resolve sensory ambiguities.

## Acknowledgments

This project has received funding from the European Union’s Horizon 2020 Framework Programme for Research and Innovation under the Specific Grant Agreement No. 720270, 785907 and 945539 (Human Brain Project SGA1, SGA2 and SGA3). We thank Frances Crabbe for help in acquiring fMRI data and Danielle Gadd for contributions to the behavioural study.

